# Monitoring Reactivation of Latent HIV by Label-Free Gradient Light Interference Microscopy

**DOI:** 10.1101/2020.12.16.423158

**Authors:** Neha Goswami, Yiyang Lu, Mikhail E. Kandel, Michael J. Fanous, Kathrin Bohn-Wippert, Erin N. Tevonian, Roy D. Dar, Gabriel Popescu

## Abstract

Latent human immunodeficiency virus (HIV) reservoirs in infected individuals present the largest barrier to a cure. The first step towards overcoming this challenge is to understand the science behind latency-reactivation interplay. Fluorescence imaging of GFP-tagged HIV has been the main method for studying reactivation of latent HIV in individually infected cells. In this paper, we report insights provided by label-free, gradient light interference microscopy (GLIM) about the changes in measures including dry mass, diameter, and dry mass density associated with infected cells that occur upon reactivation. We discovered that mean cell dry mass and mean diameter of latently infected cells treated with reactivating drug, TNF-α, are higher for cells with reactivated HIV as compared to those with latent disease. Results also indicate that cells with mean dry mass and diameter less than 10pg and 8µm, respectively, remain exclusively in the latent state. Also, cells with mean dry mass greater than 23pg and mean diameter greater than 11µm have a higher probability of reactivating. This study is significant as it presents a new label-free approach to quantify latent reactivation of a virus in single cells based on changes in cell morphology.

## Introduction

HIV is a global pandemic with over 38M infected individuals. Host cells infected by HIV may either actively produce new viral particles, or stay in a dormant state called latency, where they are capable of evading detection and drug treatments and start viral production upon perturbations. Despite the effectiveness of therapies to treat the actively replicating virus to below the limit of detection, the establishment and existence of latently infected CD4+ T-cells remains the major barrier to an HIV cure (Dahabieh et al., 2015; Richman et al., 2009; Ruelas and Greene, 2013; Siliciano and Greene, 2011). Upon removal of antiretroviral therapy (ART), the latent HIV reservoir of cells reactivates and reinitiates active replication of the virus to original viral levels in the blood. Viruses have been studied for underlying mechanisms establishing latency, and microscopy has been used to characterize gene regulation and viral decision-making at the single cell level (Bohn-Wippert et al., 2018; Hansen et al., 2018; Weinberger et al., 2008; Zeng et al., 2010). As a fluorescence-based model system for studying HIV latency may not always be available, label-free imaging methods may become more popular and in higher demand. Label-free imaging of HIV infected cells allows the observation of viral behavior without the need for a reporter-based system.

Biological samples are mostly transparent to visible light, resulting in low contrast brightfield images. The immediate solution in the early days was to resort to fluorescent staining for enhanced contrast. However, fluorescence staining is known to perturb the function of live cells (Hoebe et al., 2007). Furthermore, fluorescence imaging produces sample photobleaching and phototoxicity (Boudreau et al., 2016; Hoebe et al., 2007), thus, preventing long-term observations. The key to overcome the contrast limitation and to avoid the drawbacks of fluorescence is to use a phase-sensitive imaging modality. Phase information stems from the fluctuations in the optical pathlength introduced by the sample itself. The first such method was Zernike’s phase contrast microscopy (Zernike, 1955) in which a phase of π/2 is introduced between incident and scattered field components. Phase contrast microscopes enhance the contrast of biological samples significantly. Another advance in mitigating low contrast was the introduction of differential interference contrast microscopy (Lang, 1982) (DIC), wherein the contrast is provided by the interference of laterally sheared and orthogonally polarized fields traversing the sample. These methods fall into the category of qualitative phase imaging methods, where contrast is increased, but phase information cannot be extracted quantitatively.

In order to extract quantitative information from unlabeled specimens and take advantage of progress in light sources, modulation devices, and photodetectors, recently quantitative phase imaging (QPI) has developed very rapidly. Quantitative phase imaging (Popescu, 2011) is the field of microscopy methods that measure phase information from the specimens of interest. This capability enabled a broad range of biomedical applications (see Ref. (Park et al., 2018) for a review). QPI has been applied to investigations of cell growth (Kandel et al., 2019b; Lee et al., 2017; Mir et al., 2011; Sridharan Weaver et al., 2019), neuron dynamics (Cintora et al., 2017; Fan et al., 2017; Hu and Popescu, 2018; Hu et al., 2019a; Wang et al., 2011a), intracellular mass transport (Wang et al., 2011b), red blood cell properties (Popescu et al., 2006; Popescu et al., 2005; Popescu et al., 2007), fertility outcomes in cattle (Rubessa et al., 2020; Rubessa et al., 2019), etc. Since it provides intrinsic markers like dry mass and optical path length change, it has been successful in revealing new and crucial information in histopathology (Majeed et al., 2019; Takabayashi et al., 2018, 2019), prostate cancer (Nguyen et al., 2017b), breast cancer (Majeed et al., 2018), colorectal cancer (Kandel et al., 2017), pancreatic cancer (Fanous et al., 2020), skin cancer (Li et al., 2019), blood screenings (Mir et al., 2010), pelvic organ prolapse (Hu et al., 2019b) and kidney injury (Ban et al., 2018). Recent advances in deep learning allow the development of phase imaging with computational specificity, where synthetic fluorescence is generated computationally from label-free data (Kandel et al., 2020).

GLIM is a recently developed QPI system that combines phase shifting, common path and white light configurations, which makes it extremely stable and sensitive (Kandel et al., 2019a; Nguyen et al., 2017a). In this paper, we implemented GLIM for high-throughput well-plate scanning to study the reactivation of latent HIV in a latency model established in Jurkat cells (JLat 9.2) (Dar et al., 2014; Jordan et al., 2003). We used the intrinsic cellular dry mass from the GLIM data to study the reactivation of the latent virus. We found that activated cells exhibit statistically larger dry mass than latent ones, while the dry mass density remains comparable between the two populations.

## Results

### GLIM imaging

Reactivation of latent HIV in JLat 9.2 cells (Jordan et al., 2003) was previously studied using flow cytometry (Bohn-Wippert et al., 2017; Dar et al., 2014) and fluorescence microscopy (Bohn-Wippert et al., 2018), both using fluorescent probes expressed along with the viral genome. An aim of this study was to determine the applicability of quantitative phase imaging, specifically GLIM (Figure 1), to understand the science behind cellular dynamics upon HIV reactivation from latency. Use of GLIM allows extraction of properties that fluorescence cannot provide. Since this is the first study of HIV reactivation through GLIM, the fluorescence channel from a reporter vector of HIV was simultaneously monitored to draw the parallel.

**Figure 1.**
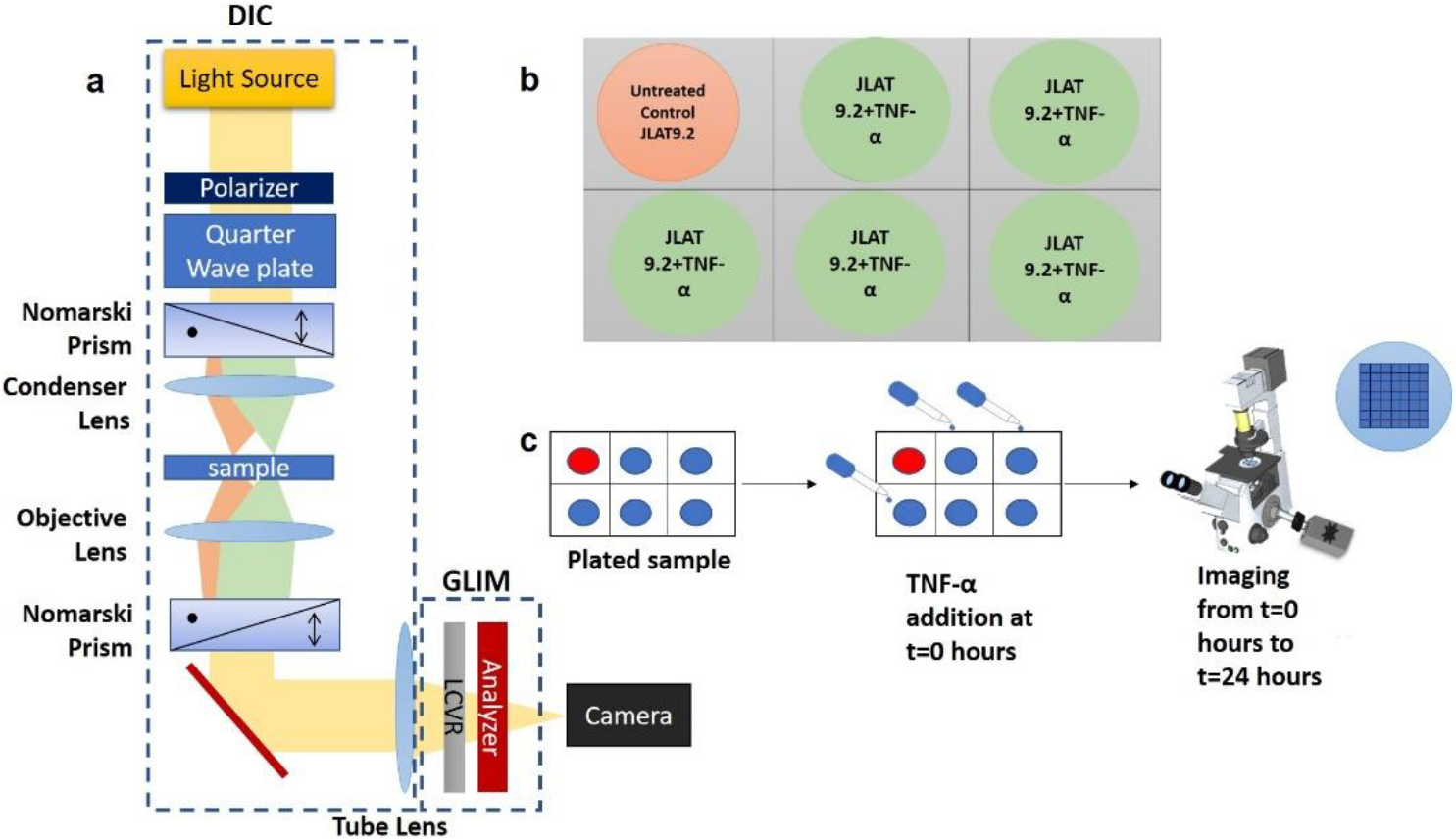
Experimental details: **(a)**. GLIM imaging setup: GLIM module is attached to a standard DIC microscope where liquid crystal variable retarder (LCVR) adds three additional phases in increments of π/2 to one of the two orthogonally sheared polarization components. **(b).** Cells were plated in a 6 well plate with untreated JLat 9.2 cells as control and replicates of JLat 9.2 treated with 10ng/mL TNF-α. **(c).** Experiment flow: TNF-α was added at t=0 hrs followed by imaging on a dual channel (GLIM+fluorescence) microscope, inset shows the 7 × 7 scanning grid per well.

JLat 9.2 cells were plated according to standard Cell-Tak protocol (see Supplementary Note 1). Tumor necrosis factor (TNF)-α, a drug that reactivates HIV from latency (Folks et al., 1989), was added at a concentration of 10ng/mL to the plated cells right before the time-lapse imaging. Figure 1a shows the GLIM imaging setup. Dual channel comprising of simultaneous GLIM and fluorescence acquisition, time-lapse imaging was performed on the sample for 24 hours. Workflow is shown in Figures 1b and 1c. Upon reactivation, cells emit green fluorescence signal due to expression of green fluorescent protein (GFP). In all repeats of the experiment, reactivation occurred at nearly 8-10 hours after the drug-addition. The percent reactivation of the whole cell population is about 20% nearly 20 hours after TNF-α addition. These are consistent with previous reports (Bohn-Wippert et al., 2018). JLat 9.2 cells are inherently in a latent state. These cells never reactivate or produce GFP signal without drug perturbations, which was utilized as the control group in all the experiments to test this claim. No fluorescence signal was observed in the control group. GLIM can provide quantitative phase information of the cells under study. Dual-channel imaging provided phase maps along with fluorescence maps for each frame. Acquired phase information is intrinsic and does not require the use of any genetic modification or staining of the sample. This enables cellular imaging in their unperturbed form. The genetic structure of the JLat 9.2 is shown in Figure 2a. Figure 2b shows the results obtained through GLIM imaging. Time-lapse imaging was performed for 24 hours showing the progression of reactivation in Figure 2c where the first frame shows the reactivation that started at nearly 9 hours after TNF-α addition (Bohn-Wippert et al., 2018).

**Figure 2.**
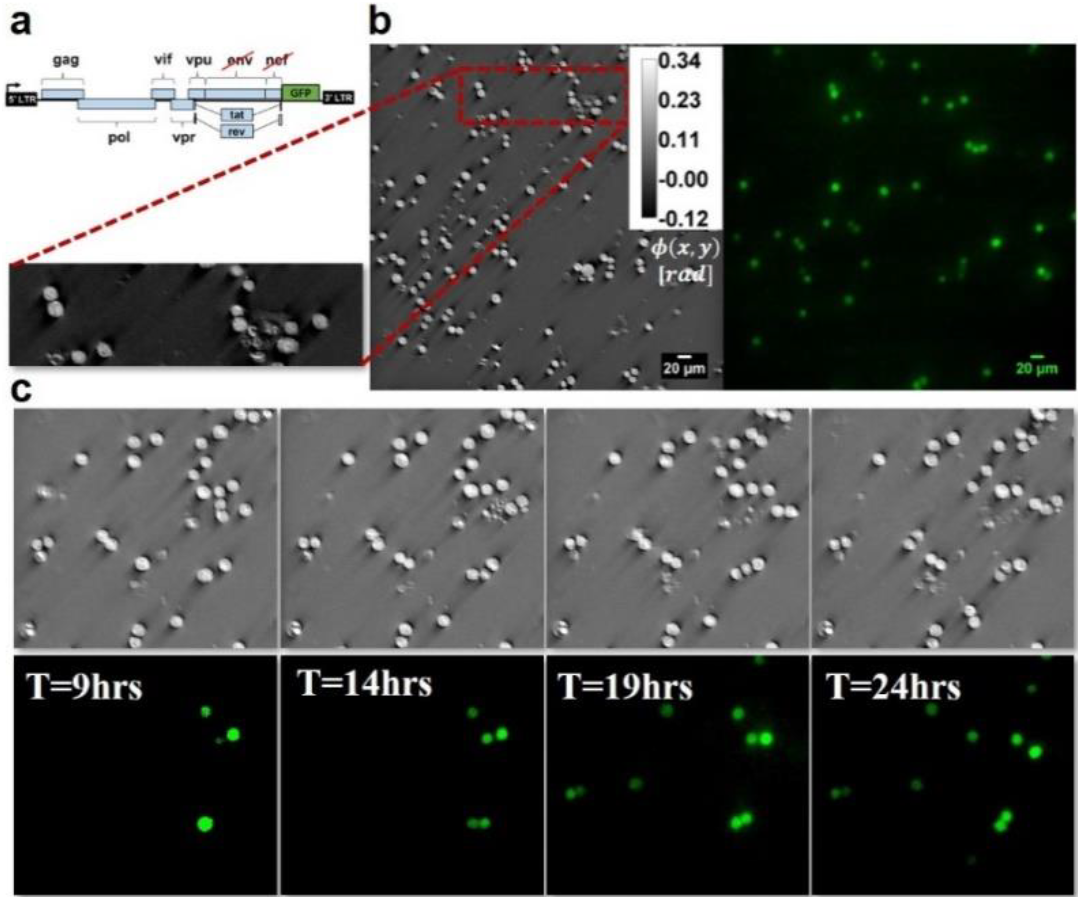
Correlative GLIM and fluorescence imaging. **(a).** JLat 9.2 is a clonal cell line derived from Jurkat T-cells, infected with a full-length HIV gene circuit, with a deletion of the *env* reading frame and a replacement of *nef* reading frame by GFP (Jordan et al., 2003) **(b).** Quantitative phase image obtained from GLIM after integrating the phase gradient map (shown in inset) and fluorescence map for same field of view. Imaging was done using DIC 20x/0.8 objective and FITC filter for fluorescence **(c).** 24-hour time-lapse imaging results-reactivation of HIV from latency was first observed around 8-10 hours after drug addition. Frames after every 5 hours interval are shown.

### Cells shift to larger diameter and higher dry mass upon reactivation

Images from both channels, GLIM and fluorescence, were registered and segmented to extract dry mass, diameter and dry mass density of individual cells. Single cell tracking was performed to quantify reactivation at the cellular level. Results of these processing steps are shown in Figure 3 (see Supplementary Note 2 and Figures S1 and S2). Note that the tracking result has memory of all the time frames, which is why there are tracks where once the cells were, detached, and left the frame. Two types of processing techniques were employed. One was the bulk processing (see Supplementary Note 2 and Figures S3 and S4), where the quantity of interest was averaged over the entire frame and in time. It provided a coarse and quick analysis. We also employed a second, more detailed analysis, in which quantities were averaged over time for each single cell. Figure 4 shows the results of single-cell tracking method. Temporal averages of cell dry mass (M) (see Methods, Eq.6) and diameter (D) show a shift towards higher end for reactivated HIV. On the other hand, shift in dry mass density (ρ) (see Methods, Eq.5) is small compared to the other two measures. These histograms also provide us the range of each measure. There were cells with diameter as low as 6µm to as high as 17µm. Similarly, the observed dry mass ranges from 8pg to 60 pg. Thus, it can be concluded that the temporal mean (signifying individual cell trend, Figure 4) display a significant difference in mean diameter and dry mass between the cells that reactivated and those that did not, with 1-way ANOVA test yielding p-values<0.001 (exact values shown in Figures 5a to 5c). These results are compatible with the changes in diameter observed in a previous study (Bohn-Wippert et al., 2018).

**Figure 3.**
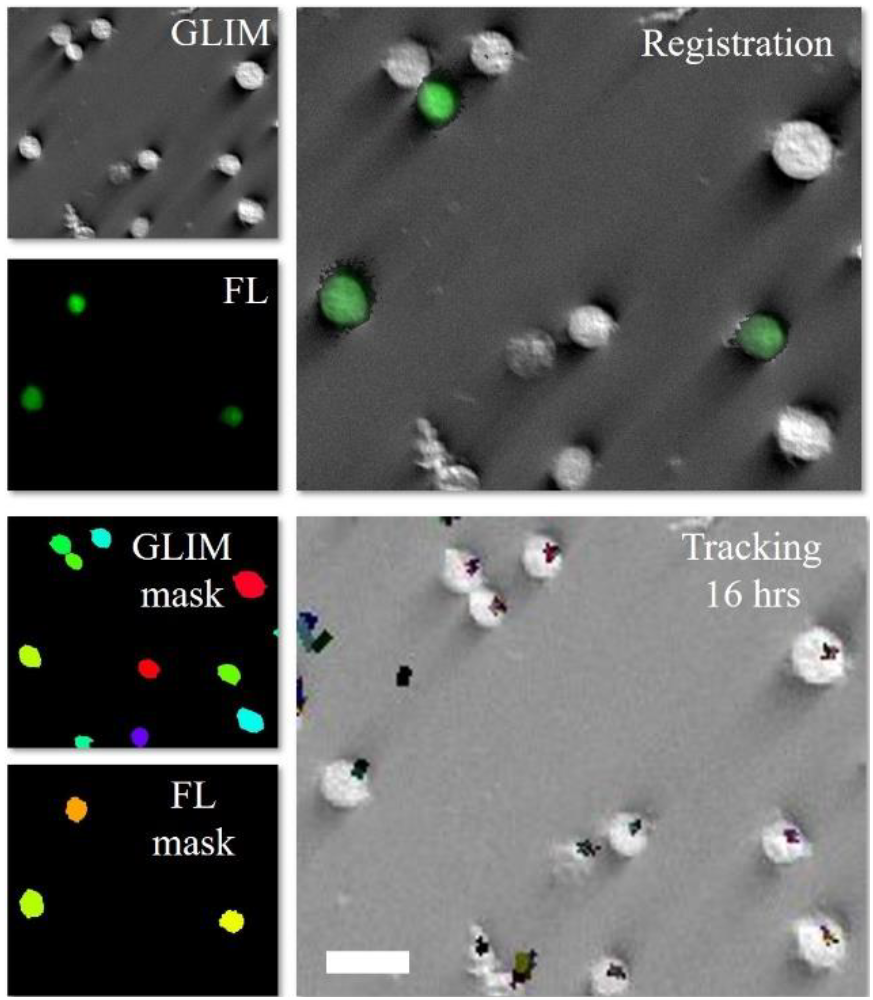
Processing workflow: GLIM and fluorescence (FL) masks were prepared from GLIM and fluorescence images (GLIM and FL) respectively. GLIM and FL were registered to overlay them perfectly. Single cell tracking was performed for 16 hours (from 9^th^ hour after reactivation to 24 hours). Single cell trajectories are shown in the tracking window. Scalebar: 20μm

**Figure 4.**
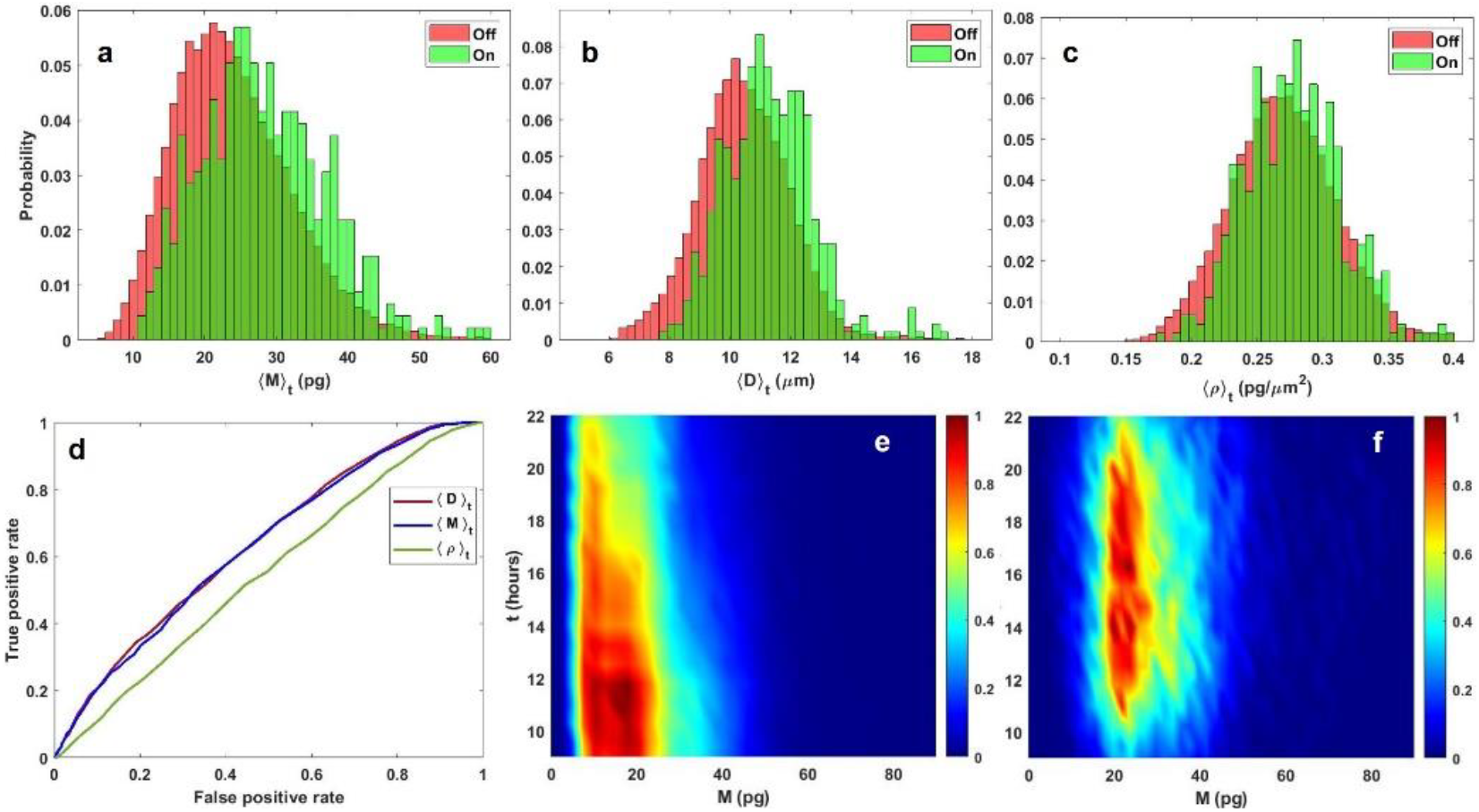
Single-cell processing result: Histogram of temporal average of **(a)**. dry mass (M), **(b).** diameter (D) and **(c).** dry mass density (ρ) respectively. The single cell processing also provides the range of values (min/max) for each quantity. For example., histogram of diameter shows that there were cells with diameter as low as 6 μm to as high as 17 μm. Legend denotes reactivated cells by ‘On’ and latent cells by ‘Off’. **(d)**. ROC curves for the three quantities. The area under curve is 0.63, 0.64 and 0.55 for dry mass (M), diameter (D) and dry mass density (ρ) respectively. Top view of dry mass histogram over time for **(e).**‘off’ cells and **(f).**‘on’ cells

**Figure 5.**
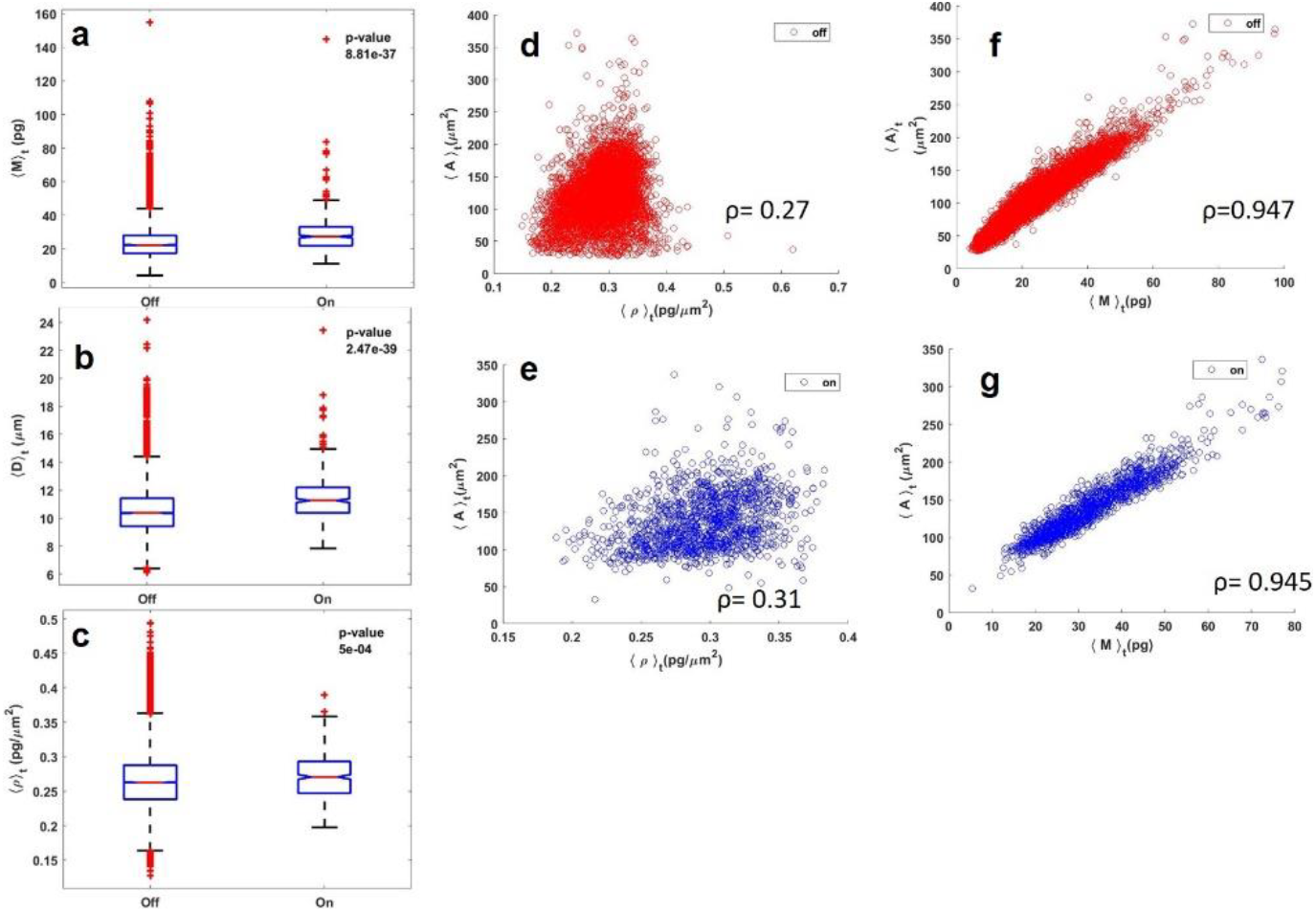
Statistical Results: 1-way ANOVA test for temporal average of **(a).** dry mass (M), p=8.81e-37, **(b).** diameter (D), p=2.47e-39 and **(c)**. dry mass density (ρ), p=5e-04. Each quantity exhibits p<0.001 denoting significance. Sample size is 24418 for ‘Off’ group and 445 for ‘On’ group. Dry mass and diameter are suitable choices for measures of reactivation because of lower p-value, indicating higher difference, as compared to dry-mass density. Area-dry-mass density scatter plot for **(d)**. ‘Off’ and **(e).** ‘On’ cells, showing no dependency. However, there is an observable dependency between area and dry-mass for both **(f).** ‘Off’ and **(g).** ‘On’ cells. Pearson’s correlation coefficients are shown on each plot.

### Cell dry mass provides a measure of reactivation but not dry mass density

Change in dry mass density was not as evident as the changes observed in diameter and dry mass. This implies that dry mass density remains a biological constant between latent and reactivated states of HIV in JLat 9.2 cells. This is further proved by ROC curves in Figure 4d, where the area under curve (AUC) for dry-mass and diameter are nearly the same (0.64 and 0.63 respectively) while the AUC for dry mass density is 0.55, which shows its limitation in differentiating between the two states. Figures 4e and 4f illustrate the top view of surface plots for dry mass of ‘Off’ and ‘On’ cells respectively over time, after reactivation. It can be seen that ‘On’ cells (Figure 4f) have larger dry mass as compared to ‘Off’ cells (Figure 4e). Another proof is the statistical result shown in Figures 5a to 5c, where the p-value of mean dry mass density is significant yet highest among the three measures. The dependency of dry mass and dry mass density on area is illustrated in Figures 5d to 5g. Dry mass density has slightly higher dependency on area for reactivated (‘On’) cells (Figure 5e) as compared to the latent (‘Off’) cells (Figure 5d). Dependence of dry mass on area is higher for both, reactivated (‘On’) cells (Figure 5g) and latent (‘Off’) cells (Figure 5f). This higher dependency of dry mass on area is expected following the relationship in Eq. (6).

### Exclusive and probable latency regime

Apart from the shift in mean dry mass and diameter, additional information was extracted from the histograms in Figure 4. It can be seen that individual cells with temporal mean dry mass less than 10pg and temporal mean diameter less than 8µm stay exclusively in the latent state. However, note that the total number of cells in these exclusive regimes is very small as compared to the total population. On the other hand, it can also be inferred that cells with mean dry mass and diameter smaller than 23pg and 11µm, respectively, have a higher probability of being latent while the ones with mean dry mass, diameter greater than these corresponding values have a higher probability of being reactivated.

### Time-trends of reactivated HIV

The reactivation of HIV in JLat 9.2 cells due to the action of TNF-α was a slow process, with the first reactivation event observed around 8-10 hours after drug addition, as shown in Figure 2c. The average ratio of reactivated and latent cells per frame is plotted with respect to time in Figure 6a. Data follows a linear trend. The range of % ‘On’ observed is consistent with reactivation previously observed for JLat 9.2 (Bohn-Wippert et al., 2017; Dar et al., 2014). Fluorescence intensity averaged over cells per time bin is shown in Figure 6b where the red curve corresponds to the mean background pixel-values of latent or non-fluorescent cells. Figure 6c shows the mean dry mass trend for reactivated cells and latent cells over time. The rate of decay of dry mass of ‘off’ cells is higher than that of ‘on’ cells.

**Figure 6.**
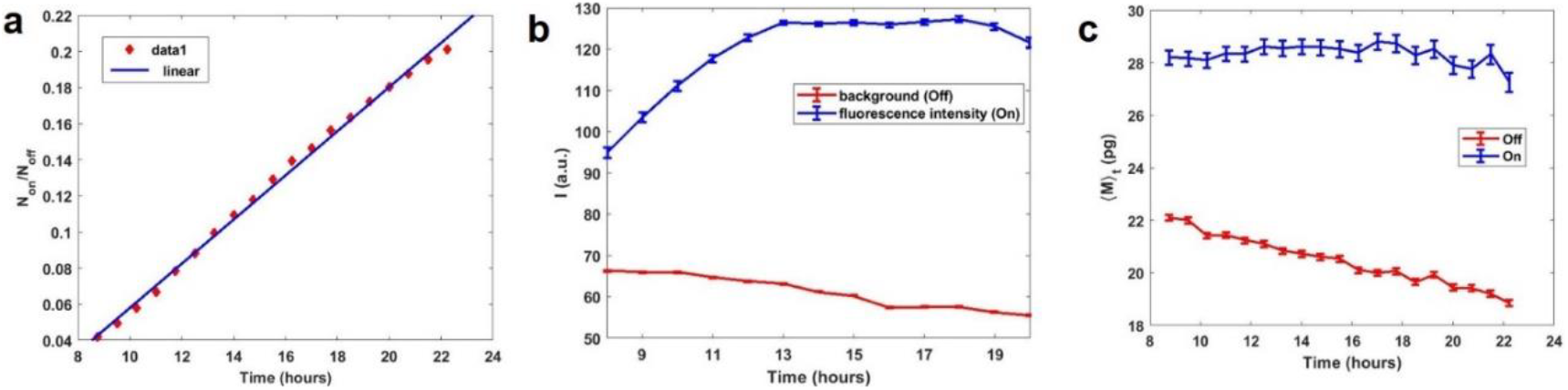
Time-trend analysis of reactivated cells: **(a).** Fraction of reactivated cells follow a growth curve, fitted with linear fit. **(b).** Mean intensity trend for reactivated, fluorescent cells (blue) and mean background pixel values for latent, non-fluorescent cells (red) **(c).** Mean dry mass time trend for reactivated (On) and latent (Off) cells. Error bars on (b) and (c) indicate standard mean error.

### Mode of reactivation

Analysis of time lapse images reveal some key modes of reactivation process. Different modes of reactivation were observed at the cellular level (Supplementary videos S1-S10). The first and most prominent mode is growth. Cells grow in size and reactivate. Another method is cell division. Reactivated cells upon division retain the reactivation state in each of the daughter cells. This explains the presence of reactivated cells in the lower end of the histograms in Figure 4.

## Discussion

We demonstrated the ability of label-free, intrinsic measures derived from GLIM to understand the differences between latent and reactivated states of HIV. Our results obtained with GLIM agree with the cell-diameter trend shown in a previous study using fluorescence imaging (Bohn-Wippert et al., 2018). In addition to the diameter, GLIM provides new intrinsic measures like dry mass and dry mass density. These measures also reflect the change in latency. Our results show that cells with higher dry mass and diameter have a higher probability of being in the reactivated state than the latent state. However, dry mass is a stronger measure as compared to the dry mass density. This result suggests that, while activation is correlated with heavier cells, this is achieved by an increase in the overall size, while the density appears to be a biological constant.

GLIM provides the ability to observe HIV reactivation events without perturbing the cell by any stains or fluorescent proteins. It has the potential to help researchers investigate various biological phenomena, based on intrinsic cell properties. This is as close as possible to track the natural cellular behavior in labs, as the chemical or genetic composition of the cell are not altered. The cells above and below dry mass cutoffs can be predicted to be latent or reactivated with a high likelihood. This study provides deeper insights into the changes incurred by cells upon HIV reactivation in terms of both size and mass. The findings of this paper are significant as they prove the potential of quantitative phase imaging in the field of virology. Our study lays the first foundation of utilizing the power of QPI in the study of HIV and potentially other viruses. HIV reactivation from latency has been shown to be cell cycle dependent (Bohn-Wippert et al., 2018), and this phenomenon will be studied through GLIM in the future. Ultimately these studies will help with the label-free evaluation of the efficiency of therapeutic strategies aiming to remove the latent cell reservoir and cure HIV (Archin et al., 2012; Deeks, 2012).

## Acknowledgements

The authors would like to acknowledge the following grants to support this study: National Institutes of Health (R01GM129709, R01CA238191), National Science Foundation (0939511, 1450962, 1353368) (awarded to G.P.). E.N.T. acknowledges support provided by the Cancer Scholars Program at UIUC. Y.L., K.B-W., and R.D.D. acknowledge support from NIH NIAID (AI120746) and NSF CAREER (1943740).

## Author contributions

G.P and R.D.D., proposed the project. Y.L., K. B-W., and E.N.T. designed and performed mammalian cell culture, protocol development. N.G. and Y.L., prepared the sample. N.G., M.E.K and M.J.F performed the dual-channel imaging. N.G did the image processing and quantitative analysis. N.G wrote the manuscript with contribution from all authors. G.P and R.D.D supervised the project and manuscript preparation.

## STAR Methods

### RESOURCE AVAILABILITY

#### Lead Contact

The lead contact for this study is Gabriel Popescu.

#### Materials Availability

We did not generate any new material in this study.

#### Data and Code Availability

All the data and code needed to reproduce the findings of this paper can be obtained from the lead contact upon a reasonable request.

### EXPERIMENTAL MODEL AND SUBJECT DETAILS

#### Cell Line

A well-established Jurkat T-cell latency model of HIV (JLat) was used in the current study (Jordan et al., 2003). In this model, T-cells are latently infected with a full-length HIV virus that has a deletion of *env* and GFP replacing the *nef* reading frame (Figure. 2a) (Jordan et al., 2003). JLat 9.2, a unique clonal latent cell population with a single integration site (Jordan et al., 2003), was chosen for the current study. JLat 9.2 cell line was obtained through the NIH AIDS Reagent Program from E. Verdin.

### METHOD DETAILS

#### Sample preparation

Cells were plated in 6-well glass bottom plates, following standard protocol (see Supplementary Note 1). Figure 1b represents the well-plate organization, one well was used as control to show the absence of GFP signal in untreated JLat 9.2 cells while the rest were plated with TNF-α treated JLat 9.2 cells.

#### GLIM

Gradient light interference microscopy (GLIM) (Kandel et al., 2019a; Nguyen et al., 2017a) is an advance microscopy method that is based on the concepts of differential interference contrast (DIC) microscopy. A low coherence source (red LED with central wavelength 690nm) is used for the microscopy, where the microscope is in the DIC configuration. The resulting spatially sheared, orthogonal polarization components are fed to a liquid crystal variable retarder (LCVR) on the output port. LCVR introduces variable phase shifts Φ_π_ in four increments of π/2 between the two sheared beams. Field at the detector plane can be described as (Kandel et al., 2019a; Nguyen et al., 2017a)

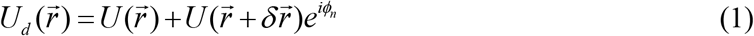

Intensity is then,

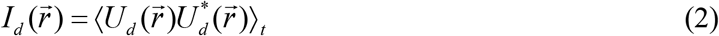

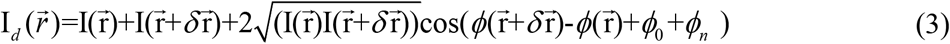

Where, ***δγ*** is the spatial displacement between two beams, *ϕ_0_* is the background phase due to the Nomarski prisms in DIC and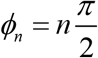, for n= 0, 1, 2 and 3 is the phase shift introduced by LCVR for phase-gradient extraction.

The quantity of interest here is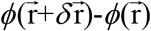, which can be expressed in terms of gradient,

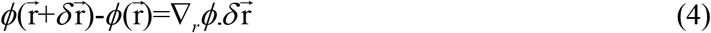

This phase-gradient is then extracted using phase-shifting interferometry algorithm (Creath, 1988; Popescu, 2011), utilizing the four intensity frames captured by varying *ϕ*_*n*_. The phase map (Figure. 2b) is obtained by integrating the gradient image.

#### Dry mass

Dry mass is a measure of non-aqueous content of the cell. Dry mass density, which is the ratio of dry mass to area, is given by (Li et al., 2019)

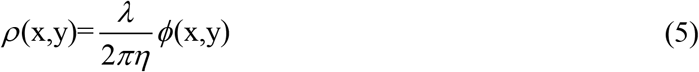

Where, *λ* is the central wavelength of light used, *η* = 0.2*mL*/*g* is the refractive index increment and *ϕ*(*x,y*) is the phase shift introduced by the sample at spatial location (x,y).

Total dry mass is calculated by taking the surface integral of *ρ* over the area of cell (Mir et al., 2011).

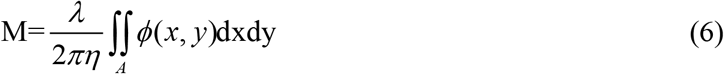

Cells were plated and TNF-α was added to all but the first well at time t=0 hour. Sample was then taken for dual-channel time lapse imaging on GLIM microscope. Each well was divided into 7 × 7 grid of frames, with each frame being imaged for both fluorescence and phase-gradient per time sampling point. Imaging was done for 24 hours. GLIM microscope is a standard DIC microscope (Zeiss Axio-observer Z1) with GLIM module (Phi Optics) attached to the side port. Cage incubator system is installed on the microscope with temperature and CO_2_ set at 37^0^ C and 5%, respectively for these experiments. Imaging was done using a 20x/0.8 DIC objective. For the fluorescence, FITC filter was used for GFP excitation. Fluorescence lamp power was kept at 50%.

#### Image Processing

After obtaining the images, they were processed using MATLAB (The MathWorks, Inc). The dual-channel images, phase and fluorescence, needed to be registered to perfectly overlay one on another. They were registered using MATLAB app, ‘Registration Estimator’. Followed by registration, images were segmented by custom build MATLAB codes, to extract binary masks for both phase and fluorescence images. These masks were used to extract underlying region properties from the images, some of which are the cell diameter and total phase of the cell from phase images and mean fluorescence intensity and identity from fluorescence images. Identity here refers to the value ‘1’ assigned to cells of the phase map that are present in the fluorescence map and ‘0’ to the rest. After feature extraction, population processing was done by averaging the cellular behavior over one frame. This was done to ascertain the population trend in latent and reactivated states of HIV. Segmentation and data extraction were followed by single cell tracking. Single cell tracking was implemented using custom build MATLAB codes. Each cell in one frame was labelled and tracked over time using Euclidean distance between centroids as a metric (see Supplementary Note 2). This was done to calculate temporal mean of the quantities of interest per cell. Single-cell analysis yielded us the range of values that were not provided by the population trend. Spatiotemporal mean is calculated for each frame and is given by

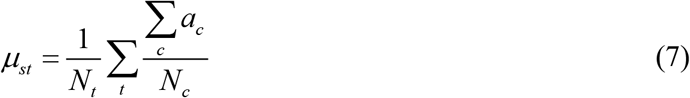

Where, *N*_*t*_ is the number of timeframes, *N*_*c*_ is the number of cells per frame and *a*_*c*_ is the quantity per cell.

Temporal mean is defined for each cell tracked over time

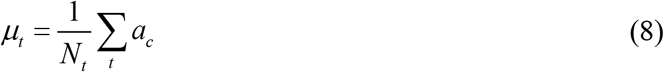

The entire processing flow is shown in Figure 3 (for more details see Supplementary Note 2 and Figures S1 and S2).

### STATISTICAL ANALYSIS

Statistical analysis of the resulting data was done using MATLAB. Since the sample size is quite large, with 445 samples in reactivated (‘On’) group and 24,418 samples in latent (‘Off’) group, normal distribution was assumed and one-way ANOVA was carried out, with default alpha 0.05. Figures 5a to 5c show that all three measures have significant difference (p<<0.001), with p-values mentioned on the bar plots, between ‘Off’ (latent) and ‘On’ (reactivated) states. To ascertain the validity of results, Kruskal-Wallis test was also carried out and it gave similar results with p-values 2.43e-32, 3.47e-38 and 5.47e-5 for dry mass, diameter and dry mass density, respectively. Kruskal-Wallis test was followed by a post hoc test on mean ranks using Scheffe’s procedure, results are shown in Figure S5.

## Supplementary Information

**Supplementary Note 1.** Cell-Tak protocol

**Supplementary Note 2.** Image processing-registration, segmentation and tracking

**Figure S1. Segmentation workflow:** Phase map was thresholded using Otsu’s method followed by hole filling and open-close morphological operations. Distance transform and extended minima functions were used to extract watershed map. Final binary mask was then obtained after watershed and filtering out irregular structures.

**Figure S2. Single cell tracking:**(a). Numbered mask for cell tracking (b). Single cell tracked over time with red line denoting centroid displacement with time (c). Single cells were tracked using Euclidean distance between centroid of the cell in successive time-frames as a metric. Each colored space represents one cell trajectory. Notice that this is a cumulative graph showing all the cells that existed in one tile. (d). Intensity trajectory over time for all cells for first five tiles of a field of view.

**Figure S3. Intensity histograms:** Histogram of fluorescence intensity for (a). Population analysis (b) Single cell analysis. There is a clear difference between the distribution of ‘Off’ and ‘On’ cells, verifying our analysis procedure.

**Figure S4 Bulk processing result:** Histogram of spatiotemporal mean for (a). dry mass, (b). diameter and (c). dry mass density. Trends match with single-cell processing results shown in main text Figure 4, showing shift in mean population dry mass and diameter towards higher end upon reactivation, while such change in dry mass density is low.

**Figure S5. Kruskal Wallis test:** Each quantity exhibits p<0.001. Sample size is 24418 in ‘Off’ group and 445 in ‘On’ group. **(a).** dry mass and **(b).** diameter are suitable choices for measures of reactivation because of lower p-value, indicating higher difference, as compared to **(c).** Dry mass density**. (d-f).** Post hoc test results: Test based on Scheffe’s procedure provides evidence of significant difference between mean ranks of all three quantities.

**Supplementary Movies S1-S10:** Movies showing the behavior of latent and reactivated cells over, ~10 hours’ time duration. Reactivated cells are labelled with green mask and latent cells are labelled with red mask.

