## Supplementary text for "Monitoring Reactivation of Latent HIV by Label-Free Gradient Light Interference Microscopy"

**Supplementary Note 1. Cell-Tak protocol**

To attach JLat cells to the bottom of imaging plates, a master mix containing Cell-Tak adhesive by Corning was first prepared. It was prepared with a mixture of sodium bicarbonate (75 g/L), sodium hydroxide (40 g/L) and Cell-Tak (2.03 g/L) at a volume ratio of 291:5:4. For each well, 187 µL of master mix was added, incubated at room temperature for more than 20 minutes, and completely dried off. Next, cells were added to the wells at a concentration of 1.5 x $10^5$ cells per well, suspended in 187 µL of DPBS. Plated cells were incubated at 5% $CO_2$, 37 °C for 30 – 60 minutes, and unadhered cells were carefully washed off with RPMI media (w/ 10% FBS and 1% pen-strep, w/o phenol red). TNF-α was diluted in RPMI media to 10 ng/ml, and 3mL of dilution was added to each well.

**Supplementary Note 2. Image processing-registration, segmentation and tracking**

Before starting the experiment, GLIM has to be calibrated following a procedure outlined in a previous work (Kandel et al., 2019a) to estimate the phase value associated with the background. The two sets of acquired data, GLIM and fluorescence were registered using Registration Estimator (a MATLAB app). This leads to a perfect overlay of phase and fluorescence images. Following the registration, segmentation was carried out using self-developed MATLAB script. Phase images were thresholded using Otsu thresholding method while fluorescence images were thresholded using adaptive thresholding. This was followed by morphological operations like hole filling, open and close etc. to maintain cell shapes. For separating conjoined cells, an algorithm based on watershed transform was employed to obtain the final mask as shown in Figure S1.

To understand the single cell behavior over time, single cell tracking was performed using MATLAB codes developed in-house. Segmented cells were numbered as shown in Figure S2a. Different parameters including cell diameter and centroid were measured for each numbered cell. Cell centroid was used as a metric to determine next neighbor of each cell in subsequent frame. The sensitivity of such detection depends on the initial estimate of velocity of cells. During the course of tracking, this initial estimate of velocity is altered to account for cells that move faster than initial estimated velocity.

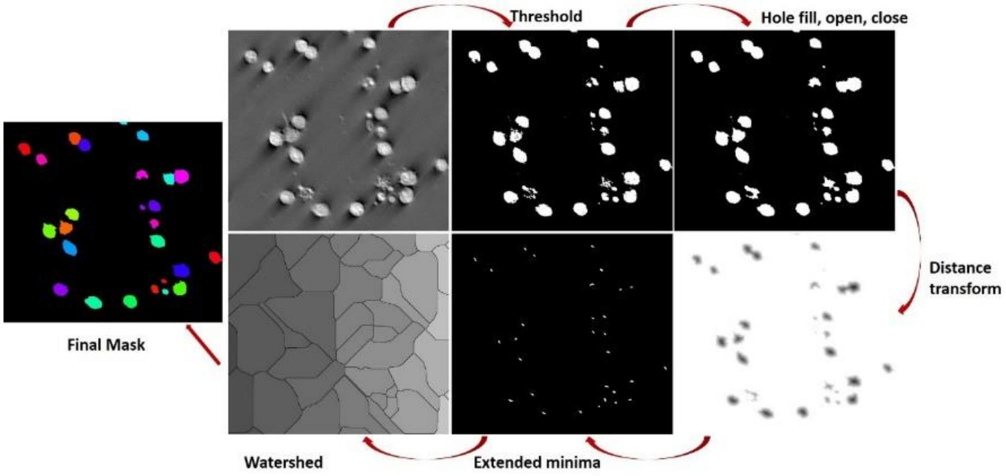


**Figure S1. Segmentation workflow:** Phase map was thresholded using Otsu's method followed by hole filling and open-close morphological operations. Distance transform and extended minima functions were used to extract watershed map. Final binary mask was then obtained after watershed and filtering out irregular structures.

Figure S2b shows the trajectories for one cell. Figure S2c shows the trajectories for all cells in one frame. It is a cumulative map showing all the trajectories of cells that existed during the time of the experiment. The mean fluorescence intensity for reactivated ('On', green) and background radiation for latent ('Off', red) cells is shown in Figure S2d. The drop in fluorescence intensity is due to photo bleaching induced by the high intensity fluorescence excitation.

### Bulk processing

Bulk/population processing is a quick alternative to single cell tracking based processing. This involves calculation of spatiotemporal means as discussed in main text, Methods section. All the cells in one frame at one time instance, signify the average behavior of one cell at that time. This is a quicker but coarse processing that provides an estimate of average population behavior.

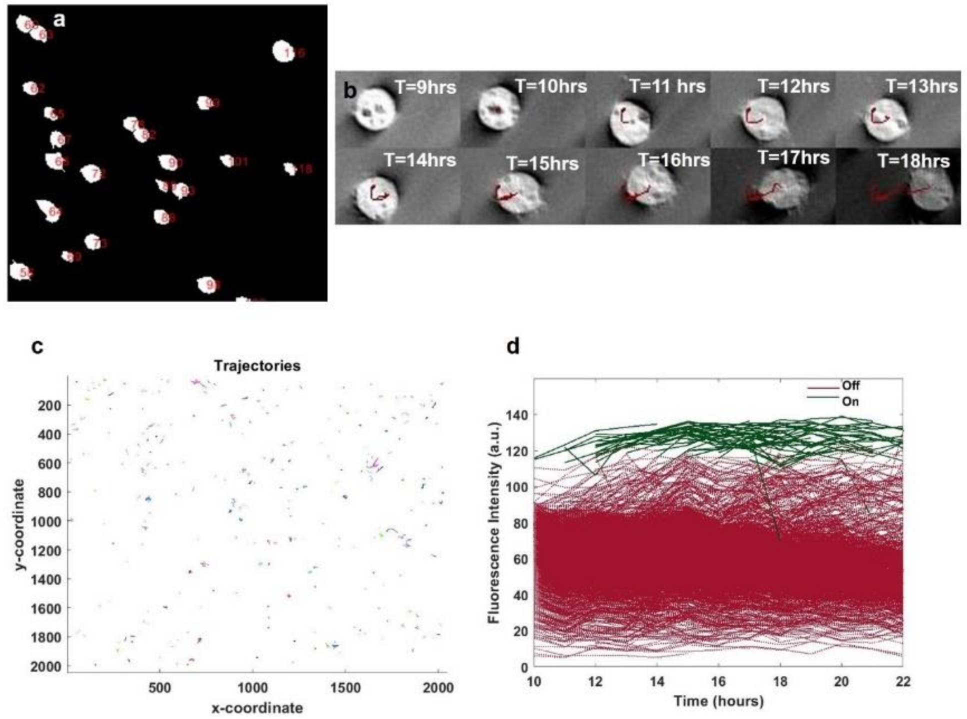


**Figure S2. Single cell tracking:** (a). Numbered mask for cell tracking (b). Single cell tracked over time with red line denoting centroid displacement with time (c). Single cells were tracked using Euclidean distance between centroid of the cell in successive time-frames as a metric. Each colored space represents one cell trajectory. Notice that this is a cumulative graph showing all the cells that existed in one tile. (d). Intensity trajectory over time for all cells for first five tiles of a field of view.

To compare both ways of processing, histograms of mean fluorescence intensity are shown in Figure S3 for bulk processing (Figure S3a) and single cell tracking (Figure S3b). Both convey similar behavior of mean but bulk processing lacks the range information conveyed by the tails in Figure S3b. However, the well-defined separation of fluorescence intensity between two groups validate our analysis procedure in both cases.

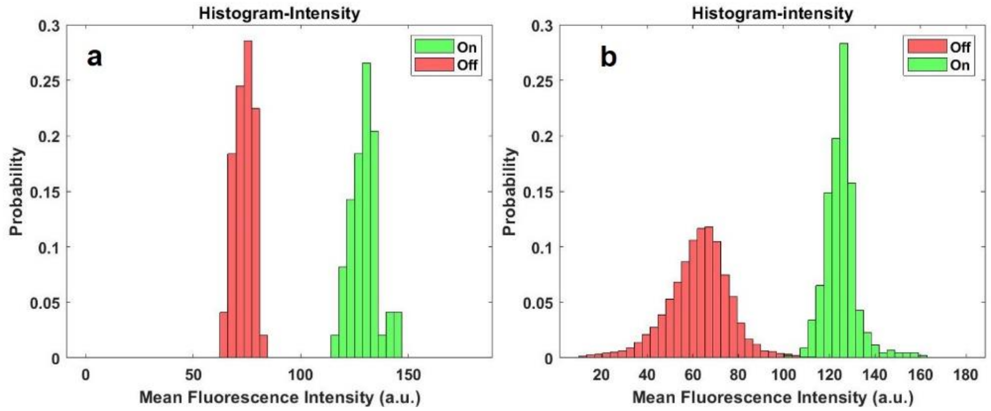


**Figure S3. Intensity histograms:** Histogram of fluorescence intensity for (a). Population analysis (b) Single cell analysis. There is a clear difference between the distribution of 'Off' and 'On' cells, verifying our analysis procedure.

The result of bulk processing for dry mass, diameter and dry mass density is as shown in Figure S4, exhibiting the shift towards higher end for dry mass (Figure S4a) and diameter (Figure S4b) in case of reactivated cells, while no such shift exists for dry mass density (Figure S4c). This finding corroborates with the single cell analysis results in main text Figure 4.

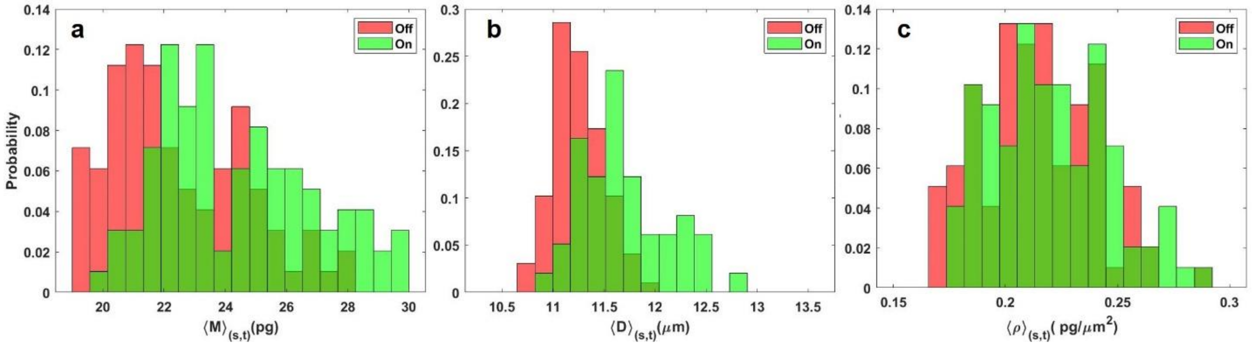


**Figure S4 Bulk processing result:** Histogram of spatiotemporal mean (a). dry mass, (b). diameter and (c). dry mass density. Trends match with single-cell processing results shown in main text Figure 4, showing shift in mean population dry mass and diameter towards higher end upon reactivation, while such change in dry mass density is low.

### Statistical Analysis: Non-parametric test

Although the large sample size in our experiment ensures normality, however, a non-parametric statistical testing of our results was done using Kruskal-Wallis test in MATLAB. The results are shown in Figure S5 which match well with the results in main text Figure 5, indicating significant separation between the dry mass and diameter for reactivated ('On') and latent ('Off') states. Dry mass density (Figure S5c) change is significant yet smaller than that in dry mass (Figure S5a) and diameter (Figure S5b). Post-hoc test based on Scheffe's procedure was carried as a follow-up test after Kruskal-Wallis test. Results agree with our observations for dry-mass, diameter and dry-mass density (Figure S5d to S5f) respectively.

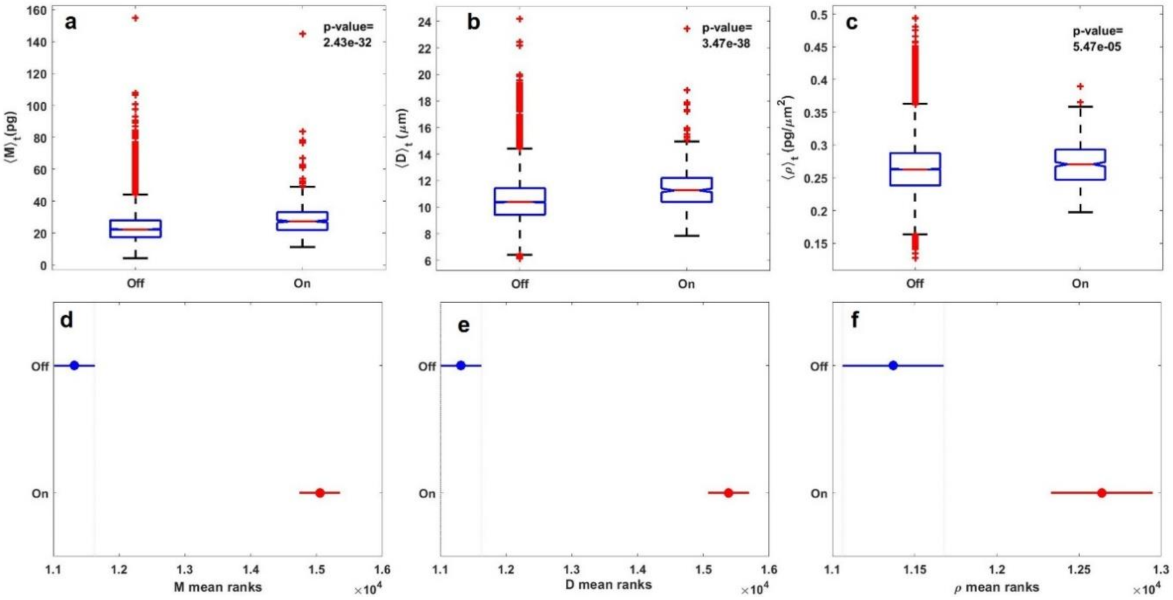


**Figure S5. Kruskal Wallis test:** Each quantity exhibits $p<0.001$. Sample size is 24418 in 'Off' group and 445 in 'On' group. **(a).** dry mass and **(b).** diameter are suitable choices for measures of reactivation because of lower p-value, indicating higher difference, as compared to **(c).** Dry mass density**. (d-f).** Post hoc test results: Test based on Scheffe's procedure provides evidence of significant difference between mean ranks of all three quantities.
